# Acute nicotine vapor normalizes sensorimotor gating and reduces locomotor activity deficits in HIV-1 transgenic rats

**DOI:** 10.1101/2024.06.18.599641

**Authors:** Neal A. Jha, Samantha M. Ayoub, M. Melissa Flesher, Kathleen Morton, Megan Sikkink, Giordano de Guglielmo, Jibran Y. Khokhar, Arpi Minassian, Arthur L. Brody, Jared W. Young

**Author notes:** Corresponding Author: Jared W. Young, Ph.D. Professor-in-Residence, UCSD Department of Psychiatry VA San Diego Healthcare System 3350 La Jolla Village Drive (116A) San Diego, CA 92161.

## Abstract

**Rationale:** Despite improved life expectancy of people with HIV (PWH), HIV-associated neurocognitive impairment (NCI) persists, alongside deficits in sensorimotor gating and neuroinflammation. PWH exhibit high smoking rates, possibly due to neuroprotective, anti-inflammatory, and cognitive-enhancing effects of nicotine, suggesting potential self-medication.

**Objectives:** Here, we tested the effects of acute nicotine vapor exposure on translatable measures of sensorimotor gating and exploratory behavior in the HIV-1 transgenic (HIV-1Tg) rat model of HIV.

**Methods:** Male and female HIV-1Tg and F344 control rats (n=57) were exposed to acute nicotine or vehicle vapor. Sensorimotor gating was assessed using prepulse inhibition (PPI) of the acoustic startle response, and exploratory behavior was evaluated using the behavioral pattern monitor (BPM).

**Results:** Vehicle-treated HIV-1Tg rats exhibited PPI deficits at low prepulse intensities compared to F344 controls, as seen previously. No PPI deficits were observed in nicotine-treated HIV-1Tg rats, however. HIV-1Tg rats were hypoactive in the BPM relative to controls, whilst nicotine vapor increased activity and exploratory behavior across genotypes. Cotinine analyses confirmed comparable levels of the primary metabolite of nicotine across genotypes.

**Conclusions:** Previous findings of PPI deficits in HIV-1Tg rats were replicated and, importantly, attenuated by acute nicotine vapor. Evidence for similar cotinine levels suggest a nicotine-specific effect in HIV-1Tg rats. HIV-1Tg rats had reduced exploratory behavior compared to controls, attenuated by acute nicotine vapor. Therefore, acute nicotine may be beneficial for remediating sensorimotor and locomotor activity deficits in PWH. Future studies should determine the long-term effects of nicotine vapor on similar HIV/NCI-relevant behaviors.

## Introduction

Of the 39 million people living with HIV (PWH) worldwide, 29.8 million are actively accessing antiretroviral therapy (ART; UNAIDS, 2023). Successful ART regimens significantly suppress HIV replication, resulting in lower viral load among PWH, enabling those with adequate ART access to live longer lives (Wada et al., 2013). The prolonged life expectancy of PWH necessitates the study of HIV as a chronic disease, requiring long-term management of the recognized cognitive and underlying neurobiological abnormalities. Despite ART, ∼50% of PWH exhibit HIV-Associated Neurocognitive Impairment (NCI), characterized by cognitive, motor, and behavioral deficits (Saylor et al., 2016). These cognitive deficits are proposed to be driven, in part, by HIV-induced neuroinflammation (Anthony et al., 2005; Peluso et al., 2013; Sreeram et al., 2022). Identifying treatments targeting such cognitive deficits and neuroinflammation would likely enhance the lives and long-term management for PWH.

PWH smoke tobacco at rates over twice that of the general population (Frazier et al., 2018) and have a higher nicotine metabolite clearance (3’-hydroxycotinine to cotinine) relative to healthy individuals (Ashare et al., 2019). These higher smoking rates and faster metabolism could relate to self-medication, especially since nicotine (the psychoactive ingredient of tobacco) has pro-cognitive effects in healthy individuals (Levin & Simon, 1998; Valentine & Sofuoglu, 2018; Wesnes & Warburton, 1983) and reduces neuroinflammation in PWH (Brody et al., 2022). However, this evidence for potential self-medication comes from associative studies, given that directionality of effects of such exposure is difficult to confirm in human studies. Additionally even if cognitively beneficial, elevated tobacco use puts PWH at higher risk of smoking-related illnesses, such as COPD (Diaz et al., 2000), which may relate to HIV protein gp120-induced mucus formation in bronchial epithelial cells (Gundavarapu et al., 2013). Further, smoking in HIV is thought to adversely affect immunological response to ART (Feldman et al., 2006). Thus, it is important to determine if nicotine directionally benefits neurocognitive function in PWH, so that future studies can determine underlying mechanisms to inform more targeted treatments.

There are a few existing clinical studies examining the neurocognitive effects of smoking in PWH, and the literature yields contradictory findings. In a group of HIV-seropositive women, a history of, but not current, smoking positively correlated with improved executive function and psychomotor speed (Wojna et al., 2007). Conversely, smoking was correlated with decreased executive function (Liang et al., 2022) and decreased psychomotor speed (Monnig et al., 2016) in PWH. However, not all clinical studies covary for Hepatitis C infection status, depressive symptoms, alcohol use, or cannabis use, each of which are elevated among PWH and independently impact cognition in HIV (Hinkin et al., 2008; Rubin & Maki, 2019; Cohen et al., 2019; Ayoub et al., 2024). Importantly, it is notoriously difficult to account for the potential impact of nicotine withdrawal, which can negatively affect cognition (Ashare et al., 2014), requiring precise timing of testing shortly after nicotine use. Furthermore, these studies are purely associative and do not infer causality, as they assess individuals who choose to use tobacco.

Among NCI-diagnosed PWH, sensorimotor gating (SG) deficits, as measured by pre- pulse inhibition (PPI), are observed (Minassian et al., 2013), though recent data indicates PPI deficits in PWH irrespective of NCI status (Walter et al., 2021). These PPI deficits are replicated in the Fischer 344 HIV-1 Transgenic (HIV-1Tg) rat model of HIV at lower pre- pulse intensities (Roberts et al., 2021a). HIV-1Tg rats are a reliable manipulation relevant to HIV, as these animals consistently express the HIV-1 proviral genome (7 of 9 proteins except for gag and pol, responsible for viral replication and reverse transcriptase) throughout the animal’s life (Reid et al., 2001). Thus, this model is ideal for studying the impact of neuroHIV in the era of ART, where the majority of PWH are virally suppressed (Hoenigl et al., 2016; Yehia et al., 2012). PPI occurs when a startle response to a high-intensity stimulus is reduced by a preceding low-intensity stimulus (Swerdlow, 2009). This attenuation is thought to relate to the pre-attentive filtering of irrelevant sensory stimuli to promote coherent thought. SG deficits are observed in various psychiatric disorders, including schizophrenia (Swerdlow et al., 2006), obsessive compulsive disorder (Ahmari et al., 2012), Huntington’s disease (Swerdlow et al., 1995), and Tourette syndrome (Castellanos et al., 1996). Interestingly, tobacco smoking enhances PPI in healthy adults (Kumari et al., 1996) and attenuates PPI deficits in people with schizophrenia (Hong et al., 2008), however the impact of such exposure on SG has not been thoroughly examined in PWH.

Preclinical studies using animal models enable controlling confounds that occur in PWH, with studies suggesting that nicotine administration, albeit via injection, has neuroprotective and pro-cognitive effects. For example, evaluation of neural structures in the HIV-1Tg rat indicated that nicotine reversed HIV-induced genomic (Cao et al., 2013) and proteomic (Nesil et al., 2015) dysregulations. Additionally, nicotine attenuated HIV-induced deficits in short-term memory and attention in the HIV-1Tg rat (Vigorito et al., 2013) and improved reinforcement learning in a different HIV model, gp120 over-expressing mice (Young et al., 2021). One study utilizing the open field test found that tobacco smoke had an anxiolytic effect selectively on HIV-1Tg rats (Royal et al., 2018), however, a follow-up study failed to reproduce this effect (Royal et al., 2022). This latter study did not quantitatively measure locomotor differences however, as their use of an open field test was to assess anxiety by time spent in the center of the field. Tracking animal movement by distance travelled and exploratory behaviors is better suited to assess locomotor activity. Additionally, in these studies (excluding Royal et al., 2018 and Royal et al., 2022), nicotine was administered via injection. It is important to model the more common practice of oral inhalation, especially with the increasing use of e-cigarettes (Tehrani et al., 2022), where pulmonary uptake of protonated (salt) nicotine formulation results in similar brain nicotine accumulation to combustible tobacco-containing cigarettes (Zuo et al., 2022). Recent studies have demonstrated the capability to conduct such vapor inhalation studies in rats, recreating self-administration and cotinine metabolite levels as in human studies (Frie et al., 2024; Frie et al., 2020; Kallupi et al., 2019; Smith et al., 2020).

To our knowledge, there are no studies examining the impact of nicotine vapor exposure on SG or locomotor activity in the context of HIV. We hypothesized that: 1) HIV-1Tg rats would exhibit PPI deficits at low prepulse levels (as seen previously); 2) acute nicotine vapor exposure would attenuate such deficits; 3) HIV-1Tg rats would exhibit abnormal locomotor and exploratory behavior; 4) acute nicotine vapor exposure would remediate such deficits; and 5) comparable levels of cotinine metabolite would be seen across genotypes after acute nicotine vapor exposure.

## Materials and Methods

### Animals

Adult (>12 weeks) male and female HIV-1Tg (n=28) and Fischer 344 (F344; n=29) control rats (65% female) were bred in-house. Animals were pair-housed in ventilated shoebox-type cages with standard environmental enrichment (plastic tube housing and nesting material; LWH: 40 x 30 x 20 cm) in a temperature-controlled (21 ± 1°C) environment on a 12-hour reversed light-dark cycle (07:00/19:00). Food and water were available *ad libitum* at all times. All testing was conducted at a minimum of 2 hours into the dark (active) portion of the rats’ photoperiod. Rats were maintained in a University of California, San Diego (UCSD) animal facility, complying with all federal and state requirements for animal care. All procedures were approved by the Institutional Animal Care and Use Committee (IACUC) at UCSD.

### Vapor Chamber Apparatus

The vapor inhalation chamber method was adapted from the OpenVape (OV) design (Frie et al., 2020). The delivery system (Fig. Ia) was constructed using a polycarbonate mouse cage (LWH: 29.8 x 18.7 x 12.7 cm) connected to a DC vacuum pump via plastic tubing. A consumer-grade SMOK e-cigarette (IVPS Technology, Shenzen, China) was fixed to the DC vacuum pump input with fitted heat shrink tubing and airtight Parafilm. The DC vacuum pumps were controlled by an Arduino microcontroller (1 per 2 cages) using the open-source code and electrical layout accessed from the Khokhar Lab website (https://www.khokharlab.com/open-source-file-downloads). The OV Arduino code remained set to pulse the e-cigarette on for 2 seconds, then rest for 4 seconds, as Frie and colleagues (2020) found this interval to maximize vapor production without overheating the e-cigarette device.

**Fig. I.**
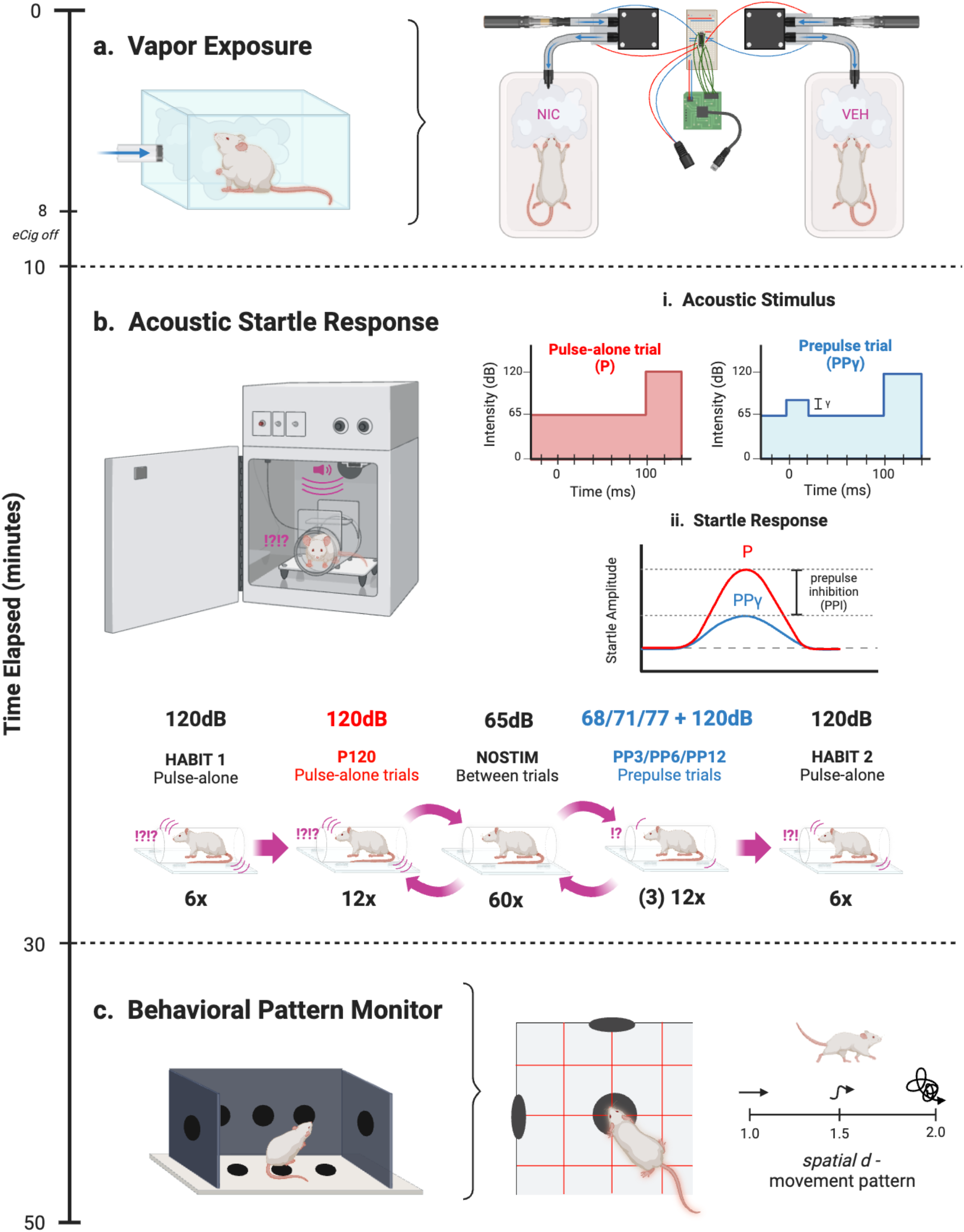
Experimental design. **a)** Rats were exposed to VEH or NIC vapor in the OV system for 10 minutes. An Arduino microcontroller controlled two miniature vacuum pumps, each connected to an eCigarette input and mouse cage output via plastic tubing. **b)** Following a 5-minute acclamation in the startle chamber, either a high-intensity 120 dB acoustic stimulus alone (P) or high-intensity stimulus 120 dB preceded by a lower-intensity stimulus (PPγ) was presented against a constant 65 dB background, where γ is the varied prepulse stimulus intensity +3, +6, or +12 dB above background (volume of 68, 71, or 77 dB, respectively). Then, ASR was assessed following the stimulus to determine PPI, the difference in startle response between P and PPγ. Testing sessions were presented in 120 trials, with no stimulus trials spaced between P and PPγ trials. HABIT trials were used to assess startle reactivity. **C)** The BPM was used to assess animal movement and exploratory behavior through holes and infrared photobeams. In addition to activity (transitions and distance travelled) and exploratory measures (nose-poking and rearing), locomotor patterns were also quantified through spatial *d*. Spatial *d* values closer to 1 indicated more linear movement, while values closer to 2 indicated more sporadic movement

### Drugs and Dosage

Nicotine (NIC) was administered by filling the e-cigarette cartridges with vape e-liquid containing protonated (salt) NIC dissolved in equal parts vegetable glycerin (VG) and propylene glycol (PG; 5% NIC, 50:50 VG/PG; VaporFi, Miami Lakes, FL, USA). For vehicle (VEH) administration, nicotine-free vape e-liquid was used (0% NIC, 50:50 VG/PG; VaporFi, Miami Lakes, FL, USA). Rats were treated with NIC or VEH via 8-minutes of on/off pump activation in the OV chamber, then remained in the chamber for an additional 2 minutes with the pump off. This drug regime was previously found to result in pharmacologically active plasma nicotine and cotinine levels in rats (Frie et al., 2020).

### Prepulse Inhibition of Acoustic Startle Response

The acoustic startle response (ASR) was measured to determine PPI using 8 startle chambers consisting of nonrestrictive cylindrical Plexiglas (8.2 cm diameter) attached to a Plexiglas platform (12.5 x 25.5 cm) with a piezoelectric accelerometer mounted underneath (Fig. Ib). Each chamber was housed in an illuminated, ventilated, sound-attenuating cabinet (SR-LAB, San Diego Instruments, San Diego, CA, USA). Acoustic stimuli were produced by a speaker mounted 24 cm above the chamber. Dynamic calibration procedures were completed prior to testing, ensuring piezoelectric accelerometers had equivalent input sensitivities and speakers had equivalent acoustic output from 60-120 dB. All animals were baseline tested and counterbalanced for drug assignment to account for underlying between-subject biological variability in PPI.

Both baseline and testing sessions consisted of various trial types following a 5-minute acclimation period: pulse-alone trials with a 40 ms-duration 120 dB pulse, prepulse+pulse trials with a varied lower-intensity 20 ms-duration pulse (68, 71, or 77 dB) preceding the 120 dB pulse by 100 ms (80 ms interstimulus interval), and no-stimulus trials including only a 65 dB background white noise. The difference in volume between the prepulse intensity and background noise was used as nomenclature for the trial type (e.g. PP3 = 68 dB – 65 dB). The trials were presented pseudorandomly for a total of 120 trials: 12 pulse-alone (P120), 12 of each varied prepulse+pulse intensity γ (PP3P120, PP6P120, PP12P120), 60 no-stimulus (NOSTIM), and 6 additional pulse-alone at the beginning (HABIT 1) and end (HABIT 2) to achieve and assess stable startle response.

NOSTIM trials were presented in an alternating order, constantly spaced between P120 and PPγP120 trials (see Fig. Ib). Startle amplitude from each trial was digitized using input from the piezoelectric accelerometers, then transmitted to a computer. ASR was defined as the arithmetic mean of accelerometer input across the recording duration of each trial. PPI at each prepulse intensity γ (PP3, PP6, and PP12) was calculated as a percent reduction in ASR from the quotient of prepulse+pulse (PPγP120) and pulse-alone (P120) trials.

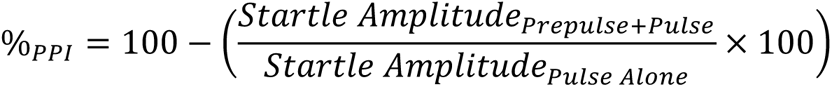

### Behavioral Pattern Monitor

The rat Behavioral Pattern Monitor (BPM; Fig. Ic), used to record animal movement and exploratory behavior, consisted of a Plexiglas chamber (LWH: 30.5 x 61 x 38 cm) illuminated by a single light source above the arena. The chamber contained 3 floor holes and 8 wall holes and was equipped with 12 x 24 infrared photobeams 1cm above the floor (2.5 cm apart) to detect hole-poking behavior. A set of 16 infrared photobeams located 2.5 cm above the floor was used to detect rearing behavior. Rat location was recorded every 100 ms with its position defined across nine unequal regions (center, 4 corners, 4 walls). At the beginning of each session, rats were placed in the center of the chamber. Activity was measured using the 12 x 24 infrared photobeams 1 cm above the floor (2.5 cm apart). Motor activity, specific exploration, and locomotor pattern were quantified by the measured variables in Table I.

**Table I.** Exploratory Behaviors Measured by the BPM. (Young et al., 2015)

| **Dimension** | **Variable** | **Description** |
| --- | --- | --- |
| Motor Activity | Counts | Total number of distinct behaviors recorded |
|  | Transitions | Number of movements from one zone to another |
|  | Distance | Total distance traveled in chamber (cm) |
| Specific Exploration | Pokes | Total number of nose pokes in any of the 11 holes |
|  | Rears | Total number of times a rat, on its hind limbs, reared into the air or against the walls |
| Locomotor Pattern | Spatial *d*<br>(Fig. 1c) | The hierarchical and geometric organization of behavior: values approaching 1 indicate linear movement, values approaching 2 indicate sporadic movement |

### Experimental Design

10-38 days after baseline PPI assessment (see timeline, Fig. I), rats were treated in the OV at least 30 minutes following transfer from the vivarium. Rats were then exposed to nicotine or vehicle vapor for 10 minutes (8-minute puff on/off time), then immediately assessed for acoustic startle response. Following the 20-minute startle session, rats were transferred to the BPM (i.e., 30 minutes post-drug exposure).

### Cotinine Metabolite Analysis

Separately from a testing day, a subset of rats (n=39) were re-exposed to vapor and following 120 minutes, trunk blood was collected via rapid decapitation. Samples of blood were centrifuged at 1000g for 20 minutes. Serum supernatants were immediately transferred to clean polypropylene tubes (0.5mL aliquot/rat) and stored at -80°C until analysis. The serum was analyzed for cotinine using a commercial competitive enzyme-linked immunosorbent assay (ELISA) kit (Calbiotech, El Cajon, CA, USA). In brief, standards and serum specimens were added in duplicate to a 96-well plate pre-coated with polyclonal cotinine antibodies. Cotinine horseradish peroxidase (HRP) enzyme conjugate was added, then incubated for 60 minutes. After wells were washed, colorless tetramethylbenzidine (TMB) substrate was added, reacting with HRP-conjugated antibodies to form a blue solution, then incubated for 10-15 minutes until a visually apparent standard gradient formed. Sulfuric acid stop solution was added and the plate was immediately visualized at 450 nm using a microplate spectrophotometer equipped with SoftMax Pro 5.4 (Molecular Devices, San Jose, CA, USA).

A calibration curve was constructed by plotting known concentrations against the inverse of corrected absorbance (signal from sample - signal from blank), as absorbance was inversely proportional to cotinine concentration in this competitive ELISA. Calibration curves were constructed in the linear range (1/Abs < 2.75) by the method of least squares and the dynamic range (1/Abs = 0.04-9.69) by a nonlinear second-order polynomial least squares procedure. Cotinine concentrations of serum specimens were calculated using the regression function within the associated range.

### Statistical Analysis

PPI data was analyzed via a repeated measures ANOVA with prepulse intensities (PP3, PP6, PP12) as within-subject factors and genotype, sex, and drug as between-subject factors where appropriate. Following an effect of prepulse intensity on PPI, separate ANOVAs were conducted on PP3, PP6, and PP12 intensity data. Given *a priori* hypotheses of 68 dB-specific genotype deficits (PP3), pairwise comparisons were used to assess differences between genotype within each drug group at PP3. ASR data was also analyzed via repeated measures ANOVA with pulse-alone trials (HABIT1, P120, HABIT2) as within-subject factors and genotype, sex, and drug as between-subject factors. BPM data and serum cotinine concentrations were analyzed via 3-factor ANOVA with genotype, sex, and drug as between-subject factors. Pairwise comparisons were used to compare groups following interactive effects. Statistical analyses were performed using IBM SPSS Statistics 28 (Armonk, NY, USA). Graphs were constructed using GraphPad Prism 9 (San Diego, CA, USA).

## Results

### Prepulse Inhibition (PPI)

HIV-1Tg rats and their F344 controls were exposed to nicotine or vehicle vapor prior to PPI assessment. Analyses revealed that sex had no effect on PPI [F(1,49)=1.69, *p*=0.2], therefore male and female data were collapsed. As expected, a main effect of prepulse intensity on PPI was observed within subjects [F(2,98)=80.079, *p*<0.001; Fig. IIa], where every louder prepulse resulted in increased PPI relative to every quieter prepulse (*p*s <0.001). Given deficits of HIV-1Tg rats at lower prepulses in prior publications (Roberts et al., 2021a), specific *a priori* hypotheses analyses were conducted at PP3.

**Fig. II.**
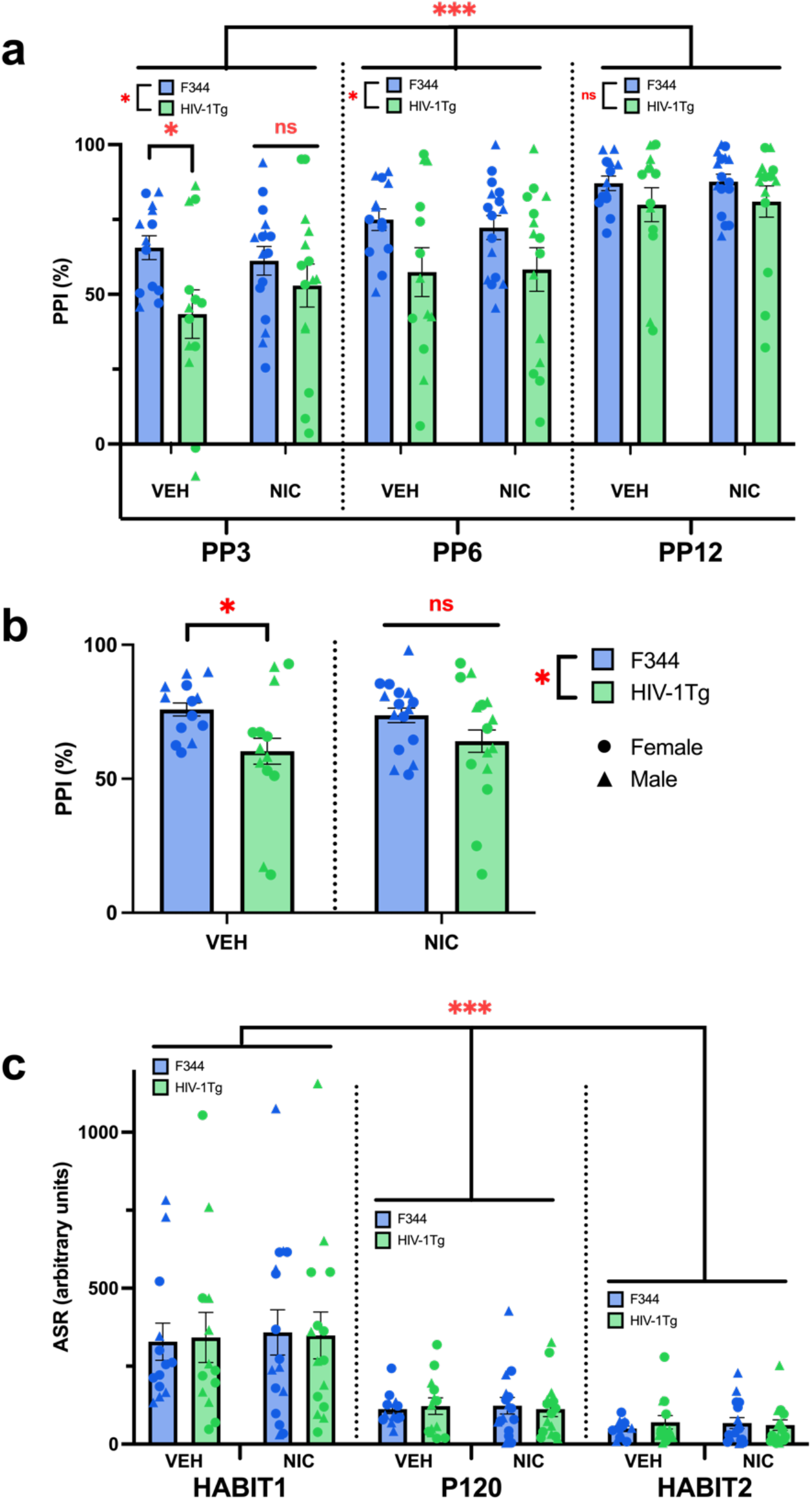
Nicotine eliminates intensity-dependent PPI deficits in HIV-1Tg rats. **a)** HIV-1Tgs had lower PPI than F344s at the lower prepulse intensities (PP3 and PP6), but not the highest (PP12). Planned comparisons at the PP3 level revealed that NIC normalized these deficits in HIV-1Tg rats. Within-subjects, louder prepulse intensities (PP3<PP6<PP12) elicited greater PPI. **b)** NIC also normalized PPI deficits when data across startle intensities were collapsed. Additionally, HIV-1Tgs exhibited lower PPI than F344s. **c)** Within-subjects, all rats had decreased startle responses to pulse-alone trials as the testing session progressed (HABIT1>P120>HABIT2). No other main effects or interactions were observed on ASR. Data presented as mean ± SEM. \**p*<0.05; \*\*\**p*<0.001; ns = not significant

Pairwise comparisons revealed that within VEH-treated rats, HIV-1Tg rats exhibited lower prepulse levels relative to their WT controls (*p*=0.019; Fig. IIa). Importantly, no genotype difference was seen in NIC-treated rats (*p*=0.329; Fig. IIa). Interestingly, this attenuating effect of NIC was also seen when prepulse intensities were collapsed, whereby the HIV-1Tg PPI deficit in VEH-treated animals (*p*=0.036; Fig. IIb) was also absent in NIC-treated animals (*p*=0.153; Fig. IIb). VEH-treated HIV-1Tg rats also exhibited reduced PPI at PP6 [F(1,53)=6.775, *p*=0.012; Fig. IIa], but not PP12 [F(1,53)=2.696, *p*=0.106; Fig. IIa]. These targeted analyses were supported by observations of an approaching intensity by genotype interaction [F(2,98)=2.434, *p*=0.093]. No other main effects or interactions were present.

### Acoustic Startle Response (ASR)

Analyses of pulse-alone trials (HABIT1, P120, HABIT2) revealed that within-subjects, a main effect of pulse-alone presentation was observed on ASR [F(2,98)=60.187, *p*<0.001; Fig. IIc), where pulse-alone trials presented later in the session elicited a lesser startle response than pulse-alone trials presented earlier in the session (*p*s <0.001). No other main effects or interactions were observed on ASR.

### Behavioral Pattern Monitor

The impact of HIV-1Tg expression in rats and response to acute nicotine vapor on locomotor and exploratory behavior was also assessed. Main effects of genotype, drug, and sex were observed on general activity levels. Specifically, HIV-1Tg rats were hypoactive, while NIC-treated and female rats were hyperactive, as measured by counts of distinct behaviors [genotype: F(1,49)=4.159, *p*=0.047; drug: F(1,49)=12.682, *p*<0.001; sex: F(1,49)=18.169, *p*<0.001; Fig. IIIa], transitions between BPM zones [genotype: F(1,49)=6.387, *p*=0.015; drug: F(1,49)=11.092, *p*=0.02; sex: F(1,49)=16.680, *p*<0.001; Fig. IIIb], and distance travelled [genotype: F(1,49)=5.930, *p*=0.019; drug: F(1,49)=12.259, *p*<0.001; sex: F(1,49)=15.132, *p*<0.001; Fig. IIIc], compared to F344, VEH-treated, and male rats, respectively. No interactions between these factors were observed for any measure.

**Fig. III.**
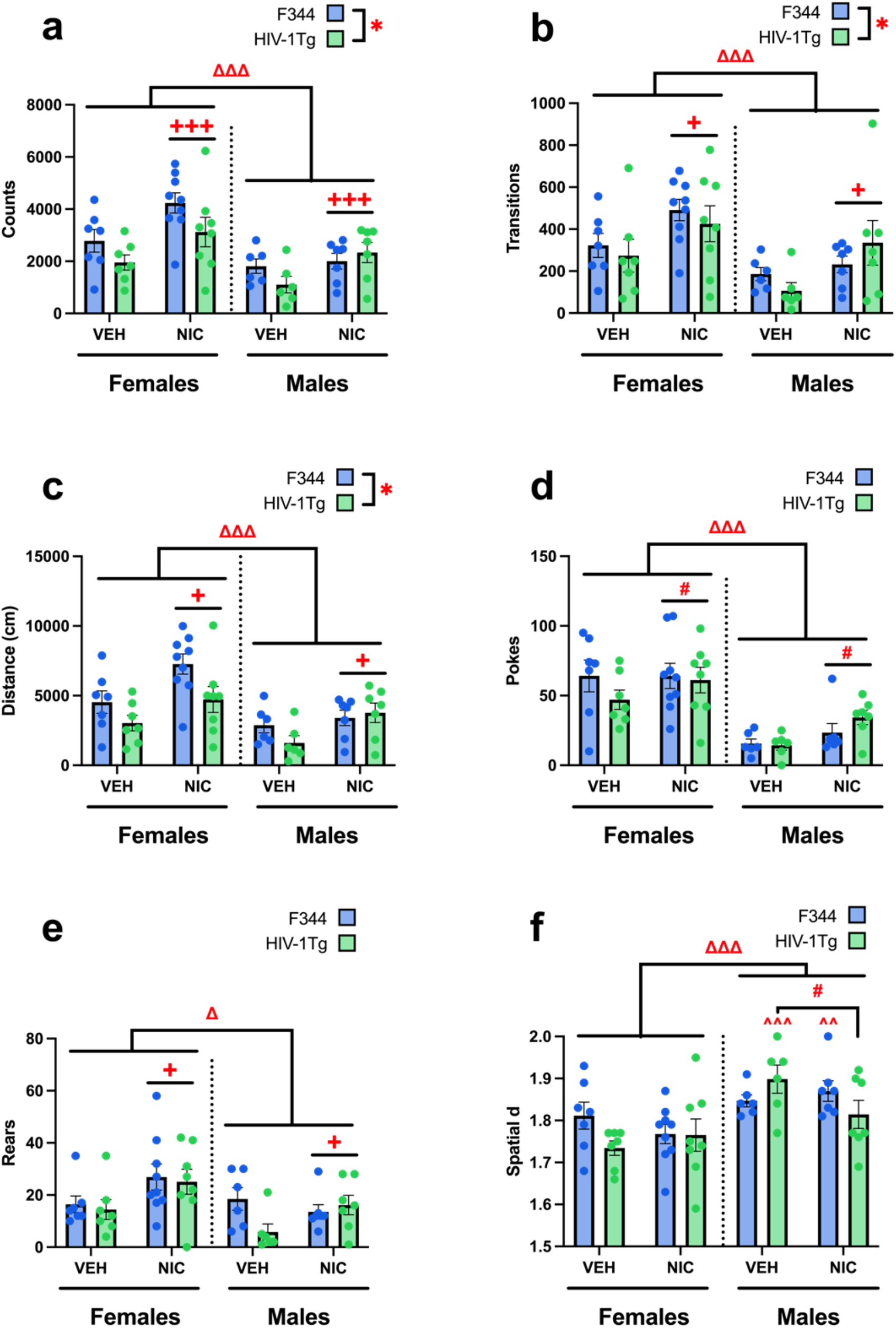
Nicotine remediates hypoactivity in HIV-1Tg rats. Males and HIV-1Tg rats had deficient motor activity, shown by less counted distinct behaviors **(a)**, transitions between BPM zones **(b)**, and total distance travelled **(c)**, which were increased by NIC across sexes and genotypes. In exploratory behavior, females made more nosepokes and NIC-treated animals trended towards more nosepokes **(d)**, and both females and NIC-treated animals had more rearings **(e)** than males and VEH-treated animals, respectively. **f)** In addition to a main sex effect of females having lower spatial *d* (more linearized locomotor pattern) than males, a drug*sex*genotype interaction revealed that NIC selectively decreased spatial *d* within HIV-1Tg males, while HIV-1Tg females were unaffected by drug. This HIV-1Tg male-specific NIC effect brought spatial *d* down to HIV-1Tg female levels, as HIV-1Tg females only had lower spatial *d* than males when given VEH. Within F344s, only NIC-treated females had lower spatial *d* than NIC-treated males. Data presented as mean ± SEM. #*p*<0.07; \**p*<0.05; \*\*\**p*<0.001, ^Δ^ sex diff. *p*<0.05; ^ΔΔΔ^ sex diff. *p*<0.001 ^+^drug diff. *p*<0.05; ^+++^drug diff. *p*<0.001; ^^within genotype-drug diff. from females *p*<0.01; ^^^within genotype-drug diff. from females *p*<0.001

**Fig. IV.**
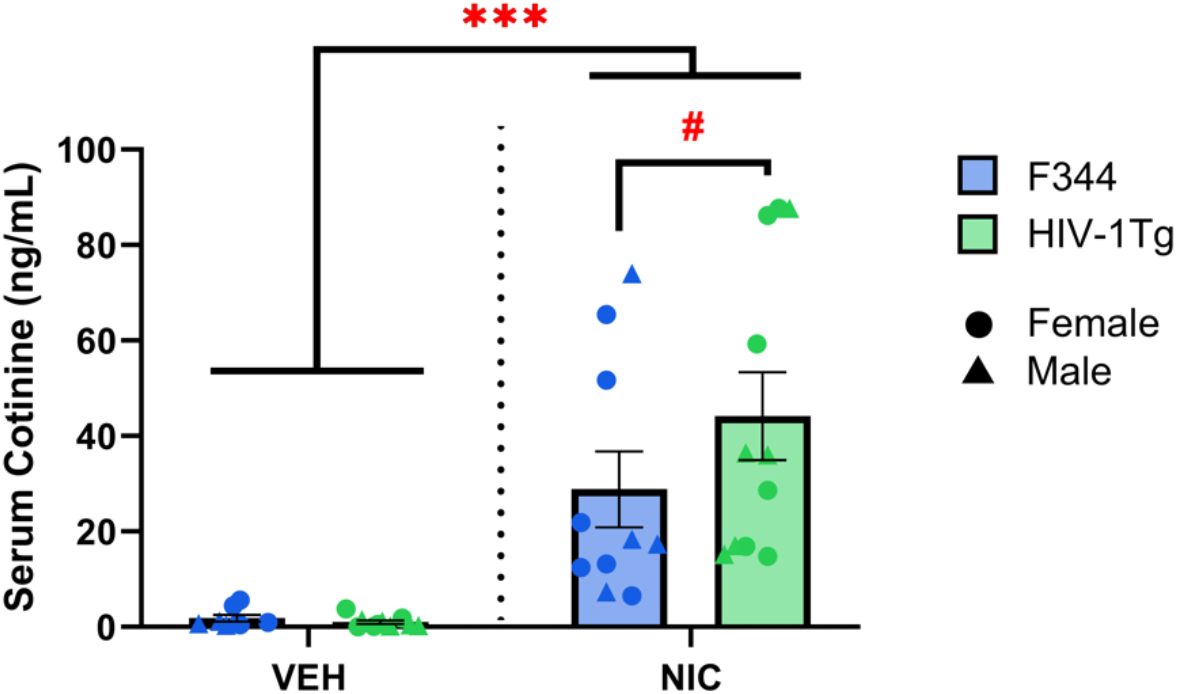
NIC-treated HIV-1Tg rats have the highest serum concentration of cotinine. Across genotypes, NIC-treated rats had higher levels of serum cotinine than VEH-treated rats. Within NIC-treated rats, the serum of HIV-1Tg rats had higher concentrations of cotinine than F344s, although this difference was only at trend levels with no interactions between genotype observed. Data presented as mean ± SEM. #p<0.1; ***p<0.001

NIC-treated and female rats exhibited increased levels of specific exploration, recorded by nosepokes [drug trend: F(1,49)=3.483, *p*=0.068; sex: F(1,49)=43.784, *p*<0.001; Fig. IIId] and rears [drug: F(1,49)=5.127, *p*=0.028; sex: F(1,49)=6.048, *p*=0.018; Fig. IIIe], than VEH-treated and male rats, respectively. No effects of genotype were observed on nosepokes or rearings. No interactions between these factors were observed for either measure.

The locomotor pattern spatial *d*—which quantifies straight line versus meandering movement—was also compared. A main sex effect was observed on spatial *d*, where females exhibited lower spatial *d* (more straight-line movement) than males (more meandering-like behavior) [sex: F(1,49)=18.735, *p*<0.001; Fig. IIIf]. Further, a genotype by drug by sex interaction was observed [F(1,49)=4.857, *p*=0.032]. Pairwise comparisons revealed that within HIV-1Tg animals, NIC tended to lower spatial *d* compared to VEH in males only (*p*=0.052), while HIV-1Tg females were unaffected (*p*=0.466). Furthermore, HIV-1Tg females exhibited lower spatial *d* than males when treated with VEH (*p*<0.001). Additionally, within F344 controls, NIC-treated females exhibited lower spatial *d* than NIC-treated males (*p*=0.009). No other main effects or interactions were observed.

## Discussion

In these studies, we demonstrated that acute nicotine vapor can normalize the PPI deficits of HIV-1Tg rat model of HIV. Consistent with previously published findings (Roberts et al., 2021a), HIV-1Tg rats exhibited deficient PPI relative to F344 controls at the lower prepulse intensities. Importantly, within the lowest prepulse intensity condition (PP3), nicotine normalized PPI in HIV-1Tg rats to levels observed in F344 control rats.

Furthermore, HIV-1Tg rats exhibited reduced locomotor activity in the BPM compared to controls, while nicotine increased this behavior in the HIV-1Tg and control rats. No main sex differences were observed during PPI testing, though females exhibited overall increased exploration and activity in every measure of the BPM. Additionally, as expected, pharmacologically relevant levels of serum cotinine were observed in nicotine-treated rats at levels comparable to prior studies in healthy rats and humans using nicotine vapor products. These results implicate the potential benefits of acute nicotine vapor on aspects of NCI in PWH, specifically revealing that acute nicotine vapor normalized SG and exploratory behavior deficits in HIV-1Tg rats.

Importantly with regards to animal modeling, here we replicated prior observations that HIV-1Tg rats exhibited impaired SG (PPI deficits) particularly at lower prepulses (Roberts et al., 2021a). Our observations extended this observed genotype effect to PP6 with overall deficits collapsed across intensities, likely due to an increased number of subjects tested (n=57) compared to the aforementioned study (n=32). Nonetheless, PPI deficits were absent in HIV-1Tg rats at the highest prepulse intensity (PP12), as previously observed. Additionally, no main effects on ASR were observed, indicating that the PPI deficits in HIV-1Tg rats were specific to SG, and not a general startle deficit. Together, these studies highlight the reproducibility of effect that PPI deficits in HIV-1Tg rats are selective to lower prepulse intensities, modeling the general PPI deficits observed in PWH (Minassian et al., 2013; Walter et al., 2021), and are seen irrespective of sex, unlike that of the gp120 (Henry et al., 2014) or iTat (Walter et al., 2021) transgenic models of HIV (deficits seen in females and males respectively). Thus, the PPI deficits of HIV-1Tg rats provide a reliable measure of deficient SG consistent with that of PWH.

Given the intensity-dependent nature of PPI deficits in HIV-1Tg rats, and our *a priori* hypotheses, we explored the effect of nicotine exposure selectively at the lowest prepulse intensity. Under PP3 conditions (68 dB prepulse), unlike after vehicle-treatment, no PPI deficits were observed in nicotine-treated HIV-1Tg rats relative to their F344 controls.

This observation suggests an attenuating effect of nicotine vapor on HIV-induced PPI deficits, consistent with previous findings of nicotine-improved PPI in people with schizophrenia (Hong et al., 2008). Interestingly, no main drug effect of nicotine on PPI was observed, despite previous demonstrations of increased PPI following acute nicotine administration, although prior rodent studies utilized injections (Acri et al., 1994; Curzon et al., 1994), and human studies were purely associative in healthy smokers (Kumari et al., 1996). Hence, acute nicotine vapor exposure specifically normalized the PPI deficits of HIV-1Tg rats at low prepulses at a dose that did not affect their control rats.

While we did not directly investigate the mechanisms underlying nicotine vapor effects on PPI in this study, it is worth speculating on the well-characterized underlying neural substrates. The nucleus accumbens (NAc) is a key regulatory region for sensorimotor gating with a prominent role of dopamine signaling (Swerdlow et al., 2001). HIV-1Tg rats exhibit reduced synaptic connectivity (McLaurin et al., 2018) and altered dopamine dynamics (Bertrand et al., 2018) in the NAc, potentially driven by the dopaminergic toxicity of HIV-1 proteins gp120 and Tat (Nath et al., 2000). Since nicotine selectively upregulates mRNA expression of *Drd5* (the gene coding for D5 dopamine receptors) in the NAc of HIV-1Tg rats but not their F344 controls (Yang et al., 2017), nicotine attenuation of PPI deficits could arise via altered dopamine dynamics. These mechanisms are likely downstream of the effects of nicotine on various nicotinic acetylcholine receptors (nAChRs), which are widely expressed in the NAc (Feduccia et al., 2012). Thus, finding the nAChRs responsible for nicotine vapor effects may yield a therapeutic target that does not have as many potential negative side-effects as nicotine, especially given that nAChRs contribute to dopamine efflux in the NAc (Maex et al., 2014). The specific mechanism through which nicotine alters PPI remains unknown, however, and further studies are certainly required to examine any cholinergic/dopaminergic interactions.

It is worth emphasizing that PPI is not a cognitive process *per se*, but rather a tool to probe the integrity of key regulatory neural substrates in the startle circuit. Further, lower PPI itself is not a diagnostic symptom of HIV or HIV-associated NCI, but rather a reflection of compromised neural substrates. For example, in decades of schizophrenia research, a causal relationship between PPI deficits and symptoms of NCI has not been found (Swerdlow et al., 2008). Additionally, in the study identifying PPI deficits in humans with HIV-associated NCI, sufficient evidence was not found to correlate SG disinhibition to general NCI (Minassian et al., 2013). Thus, our findings of nicotine-improved PPI should be interpreted simply as accompanying reports of nicotine-improved NCI in HIV rodent models (Vigorito et al., 2013; Young et al., 2021), requiring further examination of any causal relationship to other cognitive domains impacted by HIV-related NCI.

This study is the first to describe locomotor activity deficits in HIV-1Tg rats using the rodent BPM. Such studies are important given the potential for human assessment of exploration in the BPM (Minassian et al., 2022; Young et al., 2007). A previous BPM study in HIV-1Tg rats reported no genotype differences, though these rats were previously operantly trained to perform probabilistic reversal learning task, which may have skewed genotype effects on motor activity, with smaller sample sizes (Roberts et al., 2021b). Furthermore, in the current study, rats were startled in PPI chambers prior to the BPM testing session, which may have had independent stress-induced effects on locomotor behavior. Regardless, these motor activity deficits in HIV-1Tg rats (counts, transitions, distance) are consistent with human symptoms of HIV-associated NCI, particularly bradykinesia (Saylor et al., 2016). Here, acute nicotine vapor increased motor activity across genotypes, normalizing HIV-1Tg hypoactivity to levels observed in vehicle-treated F344 controls. Importantly, we replicated previous findings of increased locomotor activity in nicotine-treated rats using the OV (Frie et al., 2024) and extended these effects to HIV-1Tg rats. Together, these data support the efficacy of our vapor delivery model and suggest HIV-1tg rats do not differentially respond to acute nicotine vapor. Additionally, within male HIV-1Tg rats only, nicotine induced more linear movement. The extent to which nicotine lowered spatial *d* in HIV-1Tg males was to near-female levels however, so this interaction cannot necessarily be attributed to female HIV-1Tg insensitivity to nicotine, but perhaps a general “cutoff” of spatial *d* sensitivity to nicotine at this level. Further studies exploring dose-related and sex-specific effects are warranted.

Interestingly, nicotine vapor increased exploratory behavior (nosepokes and rears), though previous studies indicated null effects or even decreased nosepoking at higher injected doses (Geyer et al., 1986). These previous studies utilized high doses of nicotine injections, administered 10 minutes prior to testing, whereas we administered nicotine via passive vapor exposure 30 minutes prior to testing. The discrepancy in this study may be due to the different route of administration, dose, and the timeline—nicotine had longer to metabolize into a higher serum concentration of cotinine, a psychoactive compound of its own with evidence of increased locomotor activity (Wang et al., 2020). In support of this possibility, one study found that nicotine produced a biphasic effect on locomotor activity in rats, with hypoactivity during the first 10 minutes and hyperactivity at 40-50 minutes post-injection (Wiley et al., 2015). Specifically, the study by Geyer and colleagues (1986), assessed nicotine effects in the BPM during the hypoactive period of nicotine metabolism (10-30 minutes after drug), whereas our BPM assessment occurred during the likely hyperactive period (30-50 minutes after drug). Further studies are needed to dissect and compare the potential timeline and route of administration confounds between the studies.

In the BPM, females had increased motor activity, exploratory behavior, and more linear movement (lower spatial *d*) compared to males. However, sex differences are not typically observed in BPM studies cross-species (Minassian et al., 2016; Risbrough et al., 2006; Roberts et al., 2021b). Contradictory to our current findings, one recent BPM study in mice found that males had increased motor activity, exploratory behavior, and more linear movement (Ayoub et al., 2023). Although these findings are in a different rodent species, this highlights the need to consider interacting factors when interpreting sex differences, notably estrogen/testosterone pathways, each of which mediates spontaneous activity in both rats and mice (Datta et al., 2019; Miller et al., 2021; Roy & Wade, 1975; Broida & Svare, 1983). These factors certainly reinforce the inclusion of both sexes in future cross-species BPM studies to quantitatively assess distinct behaviors (as opposed to the open field test), especially given our observed interactive effect between sex, drug and genotype on spatial *d*.

Our analysis of nicotine metabolism revealed that, as expected, nicotine-treated rats had higher serum concentrations of cotinine than vehicle-treated rats. Importantly, the cotinine concentrations of nicotine-treated rats were consistent to those observed previously using an identical vaping protocol (Frie et al., 2020). The OV study used a conditioned place preference (CPP) paradigm as a proxy to assess the effectiveness of such exposure to nicotine vapor, finding that the vapor system resulted in behavioral CPP results consistent with rats receiving subcutaneous injections of 0.6 mg/kg (Torres et al., 2008). Additionally, the plasma nicotine levels associated with these cotinine levels in the OV study are consistent with plasma nicotine levels in adult cigarette smokers (Matta et al., 2007). Our replication of metabolite concentrations further validates the OV-type design as a reliable neuropharmacology tool to model eCigarette use. Additionally, since PWH exhibited faster metabolism of nicotine (Ashare et al., 2019) and higher serum concentrations of cotinine (Earla et al., 2014), we explored comparisons between drug and genotype, despite no reported genotype by drug interactions. A trend-level difference in cotinine levels was observed between nicotine-treated HIV-1Tg and control rats (*p*=0.099), but not vehicle-treated rats (*p*=0.932). Nicotine-treated HIV-1Tg rats exhibited higher levels of cotinine (mean = 44.18 ng/mL) than nicotine-treated F344 controls (mean = 28.84 ng/mL), supporting the potential for nicotine metabolic differences in HIV. This effect could also arise from blood brain barrier (BBB) abnormalities in HIV-1Tg rats, as HIV-1 protein gp120 degrades BBB permeability due to endothelial toxicity (Louboutin et al., 2010). Determining dose and duration effects of altered nicotine metabolism in HIV rats are required, particularly for future treatment development.

One limitation of our design was our choice of acute nicotine administration, whereas smoking among PWH is in the context of chronic use and addiction, wherein withdrawal effects could occur. Further, chronic assessment of PPI and BPM should be conducted, especially given findings suggesting that HIV-1Tg rats are insensitive to the locomotor-reducing effects of repeated nicotine use (Midde et al., 2011). The validity of our findings could also be enhanced by using multiple doses, or a self-administration paradigm, where rodents would voluntarily expose themselves to nicotine vapor. This approach would more accurately model human use, as shown in a study where rats self-administrating nicotine vapor had consistent blood nicotine levels with human smokers (Smith et al., 2020). Another limitation in our design was the assessment of BPM activity following acoustic startle, which may have introduced an anxiolytic confound, potentially affecting BPM data outcomes, while enabling nicotine to be further metabolized. A separate study should be conducted where rats are assessed in the BPM immediately following drug exposure to verify our hypothesis that PPI results may have been skewed by acoustic startle testing, early versus late nicotine effects, or via prolonged nicotine metabolism.

In conclusion, acute nicotine vapor exposure remediated PPI deficits and normalized the exploratory behavior of HIV-1Tg rats. PPI deficits in HIV-1Tg rats were replicated, while motor activity deficits were identified, both consistent with effects observed in PWH. These findings are important to contextualize the elevated use of nicotine among PWH and identify the potential therapeutic effects of tobacco use linked to self-medication. The translational PPI and BPM paradigms used in this study can be directly tested in humans using a similar design to validate our findings cross-species. Future studies should further explore the relationship between nicotine and HIV by investigating the underlying neurobiological mechanisms responsible for PPI modulation, modeling self-administration, and examining chronic use and withdrawal effects on these and other HIV NCI-relevant behaviors.

## Acknowledgments

This study was supported by National Institutes of Health grant R01DA051295.

The authors report no conflicts of interest.

